# The Histology of *Nanomia bijuga* (Hydrozoa: Siphonophora)

**DOI:** 10.1101/010868

**Authors:** Samuel H. Church, Stefan Siebert, Pathikrit Bhattacharyya, Casey W. Dunn

## Abstract

The siphonophore *Nanomia bijuga* is a pelagic hydrozoan (Cnidaria) with complex morphological organization. Each siphonophore is made up of many asexually produced, genetically identical zooids that are functionally specialized and morphologically distinct. These zooids predominantly arise by budding in two growth zones, and are arranged in precise patterns. This study describes the cellular anatomy of several zooid types as well as of the stem and gas-filled float, called the pneumatophore. The distribution of cellular morphologies across zooid types enhances our understanding of zooid function. The unique absorptive cells in the palpon, for example, indicate specialized intracellular digestive processing in this zooid type. Though cnidarians are usually thought of as mono-epithelial, we characterize at least two cellular populations in this species which are not connected to a basement membrane. This work provides a greater understanding of epithelial diversity within the cnidarians, and will be a foundation for future studies on *Nanomia bijuga*, including functional assays and gene expression analyses.

## Introduction

Siphonophores are pelagic hydrozoans (Cnidaria) with a highly complex development and morphological organization (Totton, 1965). Many pressing questions regarding the biology of siphonophores remain unresolved (Pugh, 1999). In particular, there have been only a few examinations of the histology of a mature siphonophore colony (e.g. Carré, 1969; Bardi and Marques, 2007) , and it remains largely unclear what cell types are present, how they are distributed across zooids, and how the tissues are organized. A better understanding of *Nanomia bijuga* histology is fundamental to questions of zooid function, colony organization, and differential gene expression.

The siphonophore *Nanomia bijuga* (delle Chiaje, 1841) begins as a single sexually produced polyp, the protozooid (Totton, 1965). The pneumatophore, a gas-filled float, forms as an invagination at the aboral or anterior end (Carré, 1969). We follow the conventions of Haddock et al. (2005) for siphonophore axes and orientation. Two distinct growth zones arise along the body column of the protozooid (Carré, 1969). At these growth zones, the protozooid body elongates and develops into the stem (Totton, 1965). New zooids are added to form a colony (Carré and Carré, 1995), with repeated zooid morphologies that are functionally and structurally specialized for particular tasks such as feeding, reproduction, and defense (Totton, 1965; Dunn and Wagner, 2006). Each zooid is a modified polyp or medusa (Totton, 1965).

The two regions that arise from these growth zones are referred to as the nectosome, which carries the pneumatophore and propulsive zooids (nectophores), which serve locomotory function, and the siphosome, which carries all other zooids (Fig. 1, Totton, 1965; Mackie et al., 1988). The siphosome contains cormidia, which are reiterated sequences of feeding zooids (gastrozooids), digestive zooids (palpons), male and female reproductive zooids (gonozooids), and protective bracts (Totton, 1965; Mackie et al., 1988). The nectosomal growth zone is located posterior to the pneumatophore, and the siphosomal growth zone is located at the anterior end of the siphosome, adjacent to the posterior of the nectosome (Totton, 1965).

**Figure 1.**
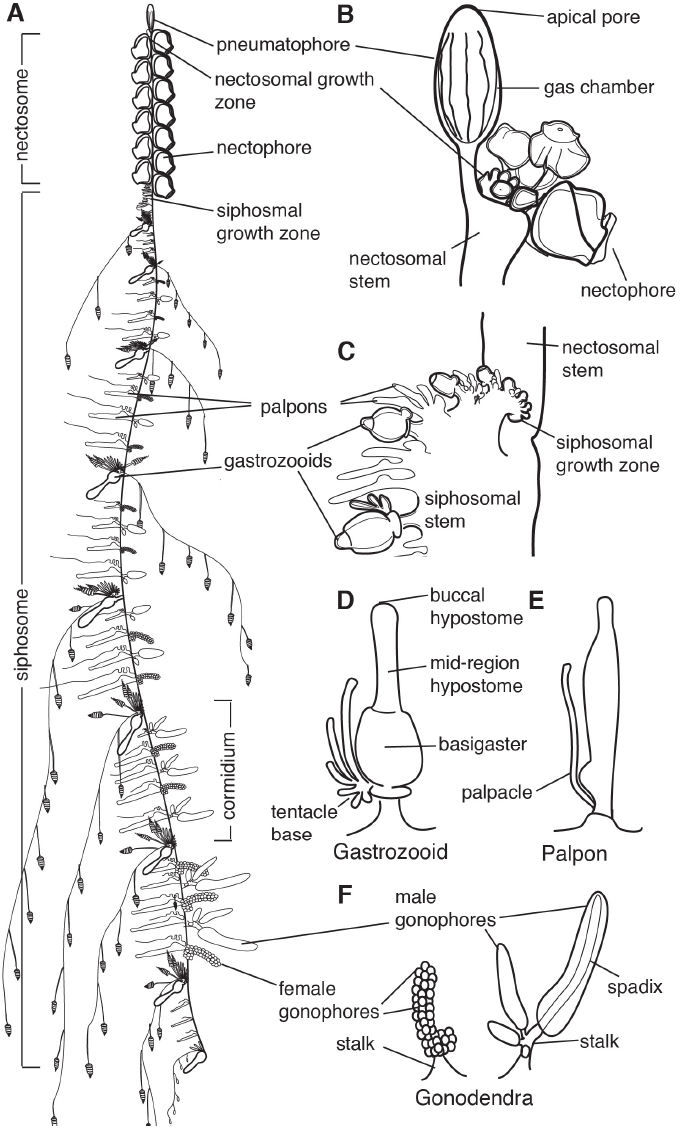
Schematic of *Nanomia bijuga* colony. Adapted from http://commons.wikimedia.org/wiki/File:Nanomia_bijuga_whole_animal_and_growth_zones.svg, which was drawn by Freya Goetz. A-C are oriented with anterior to the top and ventral to the left. D-F are oriented with anterior to the left and ventral up. (A) Overview of the mature colony. All zooids are produced from two growth zones, one at the anterior end of the siphosome, immediately posterior to the pneumatophore, and one at the anterior end of the siphosome, immediately posterior to the nectosome. Zooids are organized on the siphonosome into reiterated units known as cormidia. (B) The pneumatophore and nectosomal growth zone at the anterior of the colony, showing forming nectophores. (C) The siphosomal growth zone and budding cormidia, with gastrozooids representing cormidial boundaries. (D) Gastrozooid, connected at the base to the siphosomal stem, showing labelled regions of the mouth (buccal hypostome), the mid-region of the hypostome, and the basigaster. (E) Palpon, connected to the stem by its peduncle. (F) Female and male gonodendra. Multiple individual gonophores are borne on stalks which connect to the siphosomal stem.

Cnidarians are generally described to have diploblastic tissue layer composition; they have an outer ectoderm (also known as epidermis) and an inner endoderm (also known as a gastrodermis), separated by extracellular matrix (Thomas and Edwards, 1991). The extracellular structure on which these cells rest is known as the mesoglea and is composed of fibers in an amorphous matrix (Thomas and Edwards, 1991). Each of these cnidarian tissue layers is generally thought to be only one cell thick (mono-epithelial) (Thomas and Edwards, 1991). Stem cells, known as i-cells, were originally described in hydrozoans (Weismann, 1892). These stem cells have been shown to give rise to nematocytes, neural cells, gametes, secretory cells and, in some species, epithelial cells (Bosch and David, 1987; Plickert et al., 2012).

Several studies have touched on different aspects of the biology of *N. bijuga*, though many basic details remain unknown (Metschnikoff, 1870; Claus, 1878; Chun, 1891; Totton, 1965). Carré (1969) provided a detailed description of gametogenesis and the embryological origin of the pneumatophore, as well as descriptions of the gastrozooid. Dunn and Wagner (2006) described the organization of zooids within the colony and the budding sequence that gives rise to them. Features of the nervous system were described by Mackie (1973; 1978) and Grimmelikhuijzen et al. (1986). Siebert et al. (2011) quantified differential gene expression between zooids using RNA-seq, and demonstrated the first *in situ* mRNA hybridizations in the species. Siebert et al. (2014) described the distribution of stem cells throughout *N. bijuga*, finding that they are restricted to the growth zones, young zooids, and particular regions of some mature zooids.

This study provides a systematic investigation of the microscopic anatomy of the mature *Nanomia bijuga* colony through the use of thick sections and transmission electron microscopy, as well as a study of neural cell distribution through *in situ* hybridization. Emphasis is placed on the pneumatophore, gastrozooid, and palpon, with additional insight into the stem and gonodendra. Bracts and mature nectophores are not examined, as their enlarged gelatinous mesoglea makes them difficult to investigate with the tools used here.

## Methods

### Thick Sections and Electron Microscopy

Specimens were collected in Monterey Bay on 29 Sep 2012 via blue-water diving from a depth of 10–20 m. The specimens examined here were also used for the histological examinations of stem cells by Siebert et al. (2014). Slides with representative thick sections and three additional whole *Nanomia bijuga* specimens collected on the same dive have been deposited in the Museum of Comparative Zoology at Harvard, accession numbers IZ 50112 – IZ 50016.

Specimens were relaxed by adding isotonic 7.5 % MgCl_2_·6H_2_O in Milli-Q water at a ratio of approximately ⅓ MgCl_2_ and ⅔ FSW and fixed in 0.5 % glutaraldehyde and 4% paraformaldehyde in filtered seawater (FSW) for two minutes, transferred to 4% paraformaldehyde in FSW, and stored overnight at 4°C. After five washes with phosphate buffered saline (PBS), tissue was stored at 4°C in the presence of sodium azide [1ng/ml].

Specimens were postfixed in 2% glutaraldehyde, 4% paraformaldehyde, 100mM sucrose, and 100mM sodium cacodylate buffer. Specimens were treated with a postfix of 100mM sucrose, 100mM sodium cacodylate, and 1% osmium tetroxide overnight, after which they were washed with water and ethanol.

Tissue was processed for resin embedding according to the manufacturer’s instructions (Spurr’s Low Viscosity Resin, no. 14300). All washes and incubations were conducted at slow agitation on a rocker table.

Thick sections (0.5-0.75 μm) were prepared with glass knives, dried and counterstained for 30 seconds using toluidine blue (0.1%) in a sodium borate buffer, which stains nucleic acids blue and polysaccharides purple. Ultra thin sections (90nm) were prepared using a Diatome size 6 diamond knife (no. HI 546g). Transmission electron microscopy (TEM) images were acquired on a Philips 410 electron microscope at 80 kV. For larger structures, multiple images of a single section were aligned and blended automatically.

### In Situ Hybridization

We used tblastx (Johnson et al., 2008) against a *Nanomia bijuga* transcriptome assembly (Dunn et al., 2013) in order to identify a homolog of RFamide, a known neuropeptide in cnidarians. The *rfamide* sequence of *Polyorchis penicillatus* was used as a query (Accession number L14777). Primers were designed using Primer3 (Rozen and Skaletsky, 1999). The primers used for successful amplification were as follows: P1 - CAC ACG AAA CAG ACA TGA CAC; P2 - GTC CTT GGC TGA TTT CTC TTC. The *Nanomia bijuga rfamide (nb-rfamide*) sequence has been submitted to Genbank (Accession number KP245726). We found evidence of an additional putative *rfamide* gene in *N. bijuga* which was not characterized in this study.

*Nanomia bijuga* specimens for *in situ* hybridization were collected in Monterey Bay on July 11 and July 15, 2013 via blue-water diving from a depth of 10–20 m. After collection, specimens were kept in filtered seawater (FSW) overnight at 8°C in the dark. Specimens were transferred into a Petri dish coated with Sylgard 184 (Dow Corning Corporation) and relaxed by adding isotonic 7.5 % MgCl_2_·6H_2_O in Milli-Q water at a ratio of approximately ⅓ MgCl_2_ solution and ⅔ FSW. They were fixed in 0.5% glutaraldehyde/4% paraformaldehyde (PFA) in FSW for two minutes and incubated in 4% PFA in FSW overnight at 4°C. Specimens were then washed for three times in phosphate buffer saline and 0.1% Tween (PTw). Dehydration was performed using ethanol (EtOH) with 15 min washes in 25% EtOH/PTw, 50% EtOH/PTw, 75% EtOH/Milli-Q water, 2x 100% EtOH and then transferred to MeOH and stored at -20°C.

Dig-labeled probes were generated using Megascript T7/SP6 kits (Life Technologies). The length of the *rfamide* probe was 734 bp. Working concentration of mRNA probes were 1ng/ml. In situ hybridizations were performed according to the protocol described by Genikhovich and Technau (2009) with a few deviations. Starting at step #27, the specimens were incubated in maleic acid buffer (MAB) instead of PTw. The blocking buffer composition was MAB with 1% BSA and 25% sheep serum. Anti-Digoxigenin-AP, Fab fragments (Cat.No.11093274910, Roche Diagnostics) were used in 1:2000 dilution in blocking buffer. After antibody binding the specimens were washed in MAB instead of PBT. Once the NBT/BCIP development was stopped with water, the samples were stored overnight in 100% ethanol followed by storage in PBS. Samples were stable in PBS for many weeks provided that the medium was exchanged regularly to prevent bacterial growth. After all photo documentation was completed, specimens were stored in 4% PFA/PBS. *In situ* hybridization with multiple specimens yielded consistent results.

## Results and Discussion

### Pneumatophore

#### Background

The pneumatophore (Fig. 1A-B) is a gas filled float located at the anterior end of *Nanomia bijuga*. Carré (1969) provides a context for understanding the tissue composition of this structure and describes its development as follows. The pneumatophore arises as an invagination at the aboral end of the planula. The bilayered epithelium pushes into the gastric cavity, forming an internal mass. The ectodermal cells then divide and expand, forming two populations: a layer surrounding an internal cavity, which will become the gas chamber of the float, and a population of cells internal to this cavity which are separated from the basement membrane. The mature pneumatophore therefore has five distinct tissues (Fig. 2A). These are, from the outside inwards: external ectoderm (1) followed by its associated endoderm (2), followed by the invaginated endoderm (3), the invaginated ectoderm (4), and the population of ectodermal cells (5) within the gas chamber that are not in contact with the basement membrane. The gas chamber is surrounded by chitin and at the site of invagination there is a pore from which gas can be released. A mature pneumatophore also includes longitudinal septa that divide the gastric cavity, and which are composed of endodermal cells resting on mesoglea and connect the two gastrodermal layers. Projecting from the base of the gas chamber are blind tubes which are insinuated within the two gastrodermal layers of the septa.

**Figure 2.**
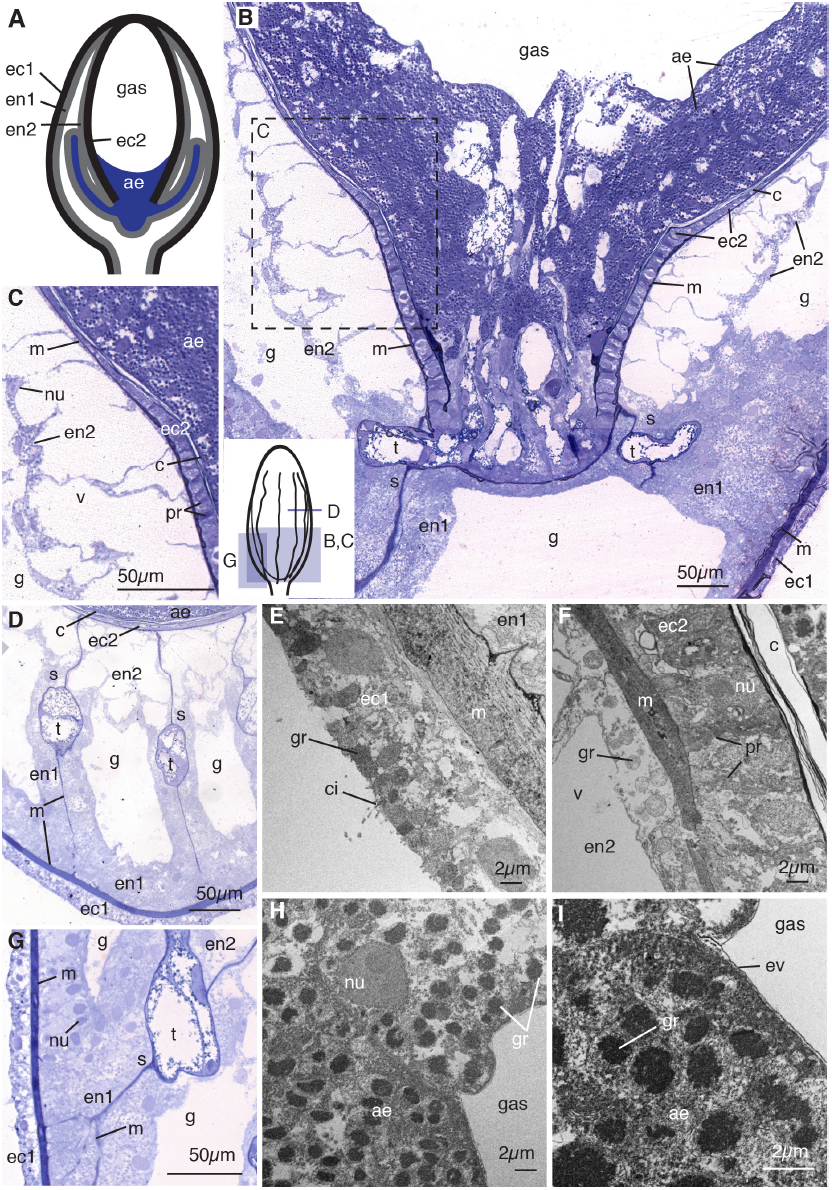
Pneumatophore, (A) Schematic of pneumatophore showing five tissue: external ectoderm (ec1),
endoderm resting on the shared mesoglea with external ectoderm (en1), invaginated endoderm (en2), invaginated ectoderm which produces chitin surrounding the gas chamber (ec2), and an ectodermally derived aeriform cell population (ae). Blind tubes connected to the base of the gas chamber are shown. (B) Longitudinal thick section of the pneumatophore base stained with toluidine blue (composite image). Small schematic at bottom left shows orientation of panels BD and G. All five tissues and multiple acellular regions are visible, including ectoderm (ec1) resting on the mesoglea (m), endodermal cells (en1) packed tightly surrounding the gastric cavity (g), invaginated endodermal cells (en2), invaginated ectodermal cells (ec2) which produce chitin (c), and the aeriform cell population (ae). At the base of the gas chamber blind tubes (t) connect and insinuate between the endodermal septa (s). (C) Enlarged image of region shown in B, showing internal endoderm, with nuclei (nu) close to the gastric membrane of the cell as well as large vacuoles (v), invaginated ectoderm with extracellular projection (pr) from mesoglea to chitin layer, and aeriform ectoderm. (D) Transverse section of the pneumatophore, showing septa composed of endodermal cells, and blind tubes. (E) Transmission electron micrograph of the transverse section of the external ectoderm (ec1), showing cilia (ci) and granules (gr). (F) Transverse electron micrograph of chitin producing cells (ec2) and chitin layer (c) and extracellular projections. (G) Thick section showing tightly packed endodermal cells, as well as the branching connections between septum mesoglea and the outermost mesoglea. (H) Transverse electron micrograph of the aeriform ectoderm cell with electron dense granules. (I) Electron micrograph of the aeriform ectoderm cells showing granules and envelope (ev) between cells and gas. Panels B-D and G are thick sections (0.5-0.75 μm) stained with toluidine blue, and panels E-F and H-I are electron micrographs (90nm).

The development and tissue composition of the mature pneumatophore has been cited as evidence in the discussion of whether the structure is medusoid or not. Most recently, Garstang (1946), Leloup (1935) and Carré (1969; 1971) established that the tissues of the pneumatophore are not homologous to those of a swimming bell, specifically because the structure is the product of invagination and not asexual budding. Additionally, the structure of the most internal population of ectodermal cells lack characteristics which would indicate homology with the entocodon of medusae (Garstang, 1946). A study by Pickwell (1964) found that the gas produced within the pneumatophore contains more than 90% carbon monoxide.

#### Results

We confirm the tissue layer arrangement of the pneumatophore as described by Carré (1969) and present additional details on the cellular features of each layer (Fig. 2). The external ectoderm contains ciliated cells with small granules close to the outer surfaces (ec1, ci, gr, Fig. 2E). The mesoglea between the external ectoderm and associated layer of endoderm is thick relative to the mesoglea in other parts of the colony (m, Fig. 2E).

The morphologies of the two endodermal cell layers are distinct (en1 and en2, Figs. 2B, D, G). The outermost endodermal cells (en1, Fig. 2B, D, G) are compact and densely arranged on the mesoglea. They are tall in shape - longer from basement membrane to the gastric surface than they are wide - and have round nuclei located at the gastric surface of the cell (nu, Fig. 2G). The second endodermal cell layer (en2, Fig. 2B-D, G), derived from the invaginated tissue during pneumatophore formation, is adjacent to the mesoglea between the gastric cavity and gas chamber and has large vacuoles (v, Fig. 2C, F). The nuclei of this endodermal cell layer are flattened and located close to the gastric surface of the cell (nu, Fig. 2C). Endodermal cells contain beta granules (gr, Fig. 2F) as defined by Fawcett (1967). The septa, which connect the outer cell layers to the inner cell layers of the gas chamber, surround the blind tubes and are composed of mesoglea lined on both sides with endodermal cells (s, Fig. 2D). The mesoglea of a septum has a single point of connection to the mesoglea of the inner endodermal cell layer but branches to form multiple connections to the outer mesoglea (s, m, Fig. 2D, G). Endodermal cells on the septa located close to the gas chamber are more similar to invaginated endodermal cells, en2, while those located closer to the external epithelial layers are more similar to en1, in that they are tightly packed and have no visible vacuoles (en1, en2, Fig. 2D, G).

A layer of chitin is located immediately interior to the layer of invaginated ectoderm (ec2, c, Figs. 2C, F). The inner chamber is surrounded entirely by chitin except at the base, where the chitin layer is not continuous (Fig. 2B). The invaginated ectoderm (ec2, Figs. 2B-C) is the only cell layer which is in contact with the chitin towards the apex of the pneumatophore, where the internal population does not extend, which supports the hypothesis that this ectoderm is responsible for the chitin secretion as suggested by Carré (1969). The ectodermal cells are wider near the base of the gas chamber and thinner toward the middle of the gas chamber (ec2, Fig. 2B). In between the individual cells of this monostratified layer are extracellular projections which span the distance between the chitin and the mesoglea; these are especially visible near the base of the gas chamber (pr, Fig. 2C, F). These connective tissues appear to attach to, but are not continuous with, the mesoglea of the invaginated ectoderm and endoderm (pr, m, Fig. 2F). The function of this layer of tissue as well as the secreted chitin shell surrounding the gas chamber is likely structural support as well as to prevent diffusion of gas out of the pneumatophore.

The cells internal to the gas chamber (ae), derived from invaginated ectodermal cells, have been referred to as aeriform (gas-producing) (Carré, 1969) or as giant cells (Garstang, 1946). The suggestion that these cells are responsible for gas production is supported by the observation that they are the only cells in direct contact with gas in the chamber, as all other cells are separated from the gas by a thick layer of chitin (ae, Fig. 2B). These cells are roughly spherical and contain many 1-2 μm granules (ae, gr, Fig. 2H-I). The population is amassed near the base of the gas chamber with no clear organization in layers (Fig. 2B). The anterior-most cells on the surface of the population have a thin layer of extracellular matrix on the exposed surface which may act as a protective envelope for the population (ev, Fig. 2I). The surface cells are separated from the mesoglea by both the chitin and multiple aeriform cells beneath them. Direct contact with mesoglea is made only by the aeriform cells at the base of the gas cavity, where there is no chitin layer, and by the cells continuous with the aeriform tissue present in the blind tubes connected to the chamber base (Fig. 2B, D).

### Gastrozooid

#### Background

The gastrozooid is a polyp that is specialized for feeding. It has two distinct regions: the oral hypostome and the aboral basigaster (Carré, 1969). The hypostome is further divided into the buccal region and the mid region, and has a thickened endoderm and thin ectoderm (Carré, 1969). The gastrozooid is capable of spreading around prey during feeding (Mackie et al., 1988). Carré and Carré (1995) described the endodermal cells in the buccal region of the hypostome as epithelio-muscular-glandular in type. In contrast to the hypostome, the basigaster has a relatively thick ectoderm that is a site of nematocyst production (Carré and Carré, 1995). This region has been referred to as the cnidogenic swelling, and has been studied for insight into the development of the stinging capsule cells (Carré and Carré, 1995). The gastrozooid tentacle is attached to the base of the basigaster on the anterior side.

#### Results

The endodermal cells of the hypostome are glandular (en, Fig. 3A-F) and club shaped, with the head of the club adjacent to the gastric cavity where secretory vesicles are released (Fig. 3F). In this study we find that the hypostome endoderm contains three types of gland cells (gc1, gc2, and gc3, Fig. 3D-F), similar to those found in other hydrozoans (Thomas and Edwards, 1991).

**Figure 3.**
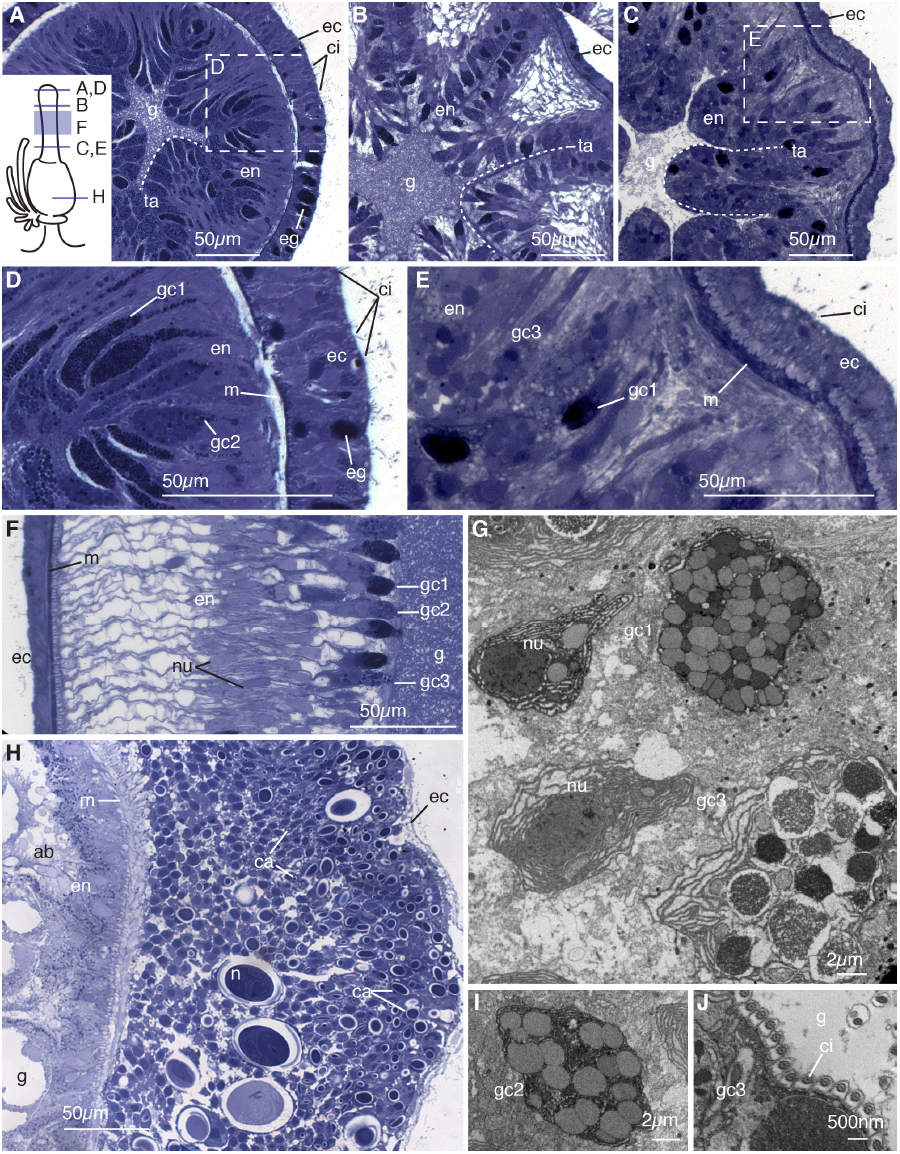
Gastrozooid. (A) Transverse section of mouth region of hypostome. Schematic in bottom right shows orientation of panels A-E and H. Visible are ectoderm cells (ec) with cilia (ci), ectodermal gland cells (eg), and club shaped gland cells of the endoderm (en), which line the gastric cavity (g) in folds known as taeniolae (ta) represented with a dashed line. (B) Transverse section of lower buccal region of hypostome, showing compact taeniolae of endodermal tissue. (C) Transverse section of mid-region of hypostome showing taeniolae of endodermal tissue. (D) Enlarged image of mouth region from A, showing ectoderm, mesoglea (m), and endoderm. Two types of endodermal gland cells are visible, one with more darkly stained granule (gc1) and one with larger granules (gc2). Ectodermal cilia and ectodermal gland cells are also visible. (E) Enlarged image of mid-region of hypostome from C, showing third type of gland cell (gc3) which contains granules which stain lightly and are much larger than those of gc2. The cilia on the ectoderm are less dense in the mid region of the hypostome. (F) Longitudinal section of the hypostome showing extended gland cells of all three types, with visible nuclei (nu) . (G) Transverse electron micrograph of the mid region of the hypostome, showing gland cells types gc1 and gc3. (H) Transverse thick section of the basigaster region. Nematocysts (n) at multiple stages of formation are visible, with undifferentiated cells near the mesoglea and capsules (ca) present in cells in middle of the cell population and those near the ectoderm. Only the undifferentiated cells nearest the gastric cavity are in contact with mesoglea; the ectodermal cell layer and all other capsule bearing cells do not appear to have a connection to the basement membrane. Absorptive cells (ab) are present endodermally. (I) Transverse electron micrograph of the hypostome showing gland cell type gc2. (J) Transverse electron micrograph of the mid-region of the hypostome, showing the ciliated surface of the gland cells, with cilia oriented perpendicular to gastric surface. Panels A-F and H are thick sections (0.5-0.75 μm) stained with toluidine blue, and panels G and I-J are electron micrographs (90nm).

In the buccal region, there are two types of gland cells: those with smaller granules which here stain darkly (gc1, Fig. 3D-G), and those with larger, more lightly stained granules (gc2, Fig. 3D, F, I). The cells with smaller, tightly packed granules (gc1) are similar to the granular mucous cells described in other hydrozoans (Siebert et al., 2008) and likely correspond to the spherical hypostomal gland cells described by Carré (1969). The buccal cells with larger, more lightly stained granules are the spumous cells (Carré, 1969). In the mid-region of the hypostome, a third gland cell type is present (gc3, Fig. 3E-G) which contains larger granules of variable stain affinity. These appear to correspond with the zymogen gland cells described in other hydrozoans (Siebert et al., 2008) as well as the gastric spherical cells described by Carré (1969). The buccal region of the hypostome contains a number of folds, referred to as taeniolae in other hydrozoans (Campbell, 1967), that presumably afford the zooid the ability to stretch around larger prey (ta, Fig. 3A-C). The taeniolae become more defined toward the mid region (ta, Fig. 3B), and more numerous and less regularly arranged further down the zooid (ta, Fig. 3C). Externally, the taeniolae appear as longitudinal stripes. They extend almost to the center of the gastric cavity (ta, Fig. 3A-C). The gastrodermal surface is densely ciliated (ci, Fig. 3J).

The ectoderm of the hypostome contains simple epithelial cells as well as gland cells (ec and eg, Fig. 3A, D). The gland cells are only present in the buccal region and also contain small, darkly stained granules (eg, Fig. 3A, D). Ectodermal ciliated cells are present more densely toward the mouth (ci, Fig. 3A, D-E). Nematocysts were not observed in the hypostome of the gastrozooid.

The ectoderm of the basigaster is roughly 25 cell diameters thick from mesoglea to surface and is the site of nematogenesis (Fig. 3H). The cells of the nematoblast population reside between the mesoglea and a very thin monolayer of ectodermal cells (ec, Fig. 3H). Within this population, the cells near the mesoglea have round nuclei that occupy most of the cellular space with developing capsules mostly absent (Fig. 3H). The cells nearer to the surface of the ectoderm have capsules at different stages of maturation (ca, Fig. 3H). Interspersed throughout these ectodermal cells are fully mature nematocysts, which measure 10-25μm (n, Fig. 3H). Large capsules are present near the base of the gastrozooid, away from likely points of contact with prey, suggesting migration to other regions of deployment within the colony. The endoderm of the basigaster consists of absorptive cells with large vacuoles and small granules (ab, Fig. 3H).

### Palpon

#### Background

The palpon is a polyp and is thought to be homologous to the gastrozooid but having lost the ability to feed (Totton, 1965). Recent findings using molecular markers and histology are in support of this hypothesis (Siebert et al., 2014). The palpon attaches to the stem via a peduncle and has a single palpacle homologous to the tentacle of the gastrozooid. The palpon has been described as an accessory of the digestive system and can be seen inflating and deflating with gastric fluid (Mackie et al., 1988).

#### Results

The gastrodermal surface of the palpon is populated by a unique cell type, previously described in *Apolemia* as ciliated funnel cells (Willem, 1894). These cells are ovoid and have at their apex a tuft of cellular projection (fc, Fig. 4A-B, D). Electron micrographs reveal that in *Nanomia bijuga* these projections are microvilli and not cilia (mi, Fig. 4E, G), therefore we refer to these cells simply as funnel cells. The microvilli are arranged in rows at the apex of the cells (mi, Fig. 4G). Inside the funnel cells are large vacuoles containing visible particulate matter (v, Fig. 4E). Large absorptive cells, densely packed with vacuoles, are also present in the endoderm (ab, Fig. 4A-B, D). The presence of these two cell types suggests that a key function of the palpon gastrodermis is particle capture and intracellular digestion. Many cells of the endoderm have two distinct nuclei which do not appear to be in the process of nuclear division (nu, Fig. 4B), and additionally well developed endoplasmic reticula are visible in endodermal cells (er, Fig. 4G). The implication of these two features for palpon function remains unknown.

**Figure 4.**
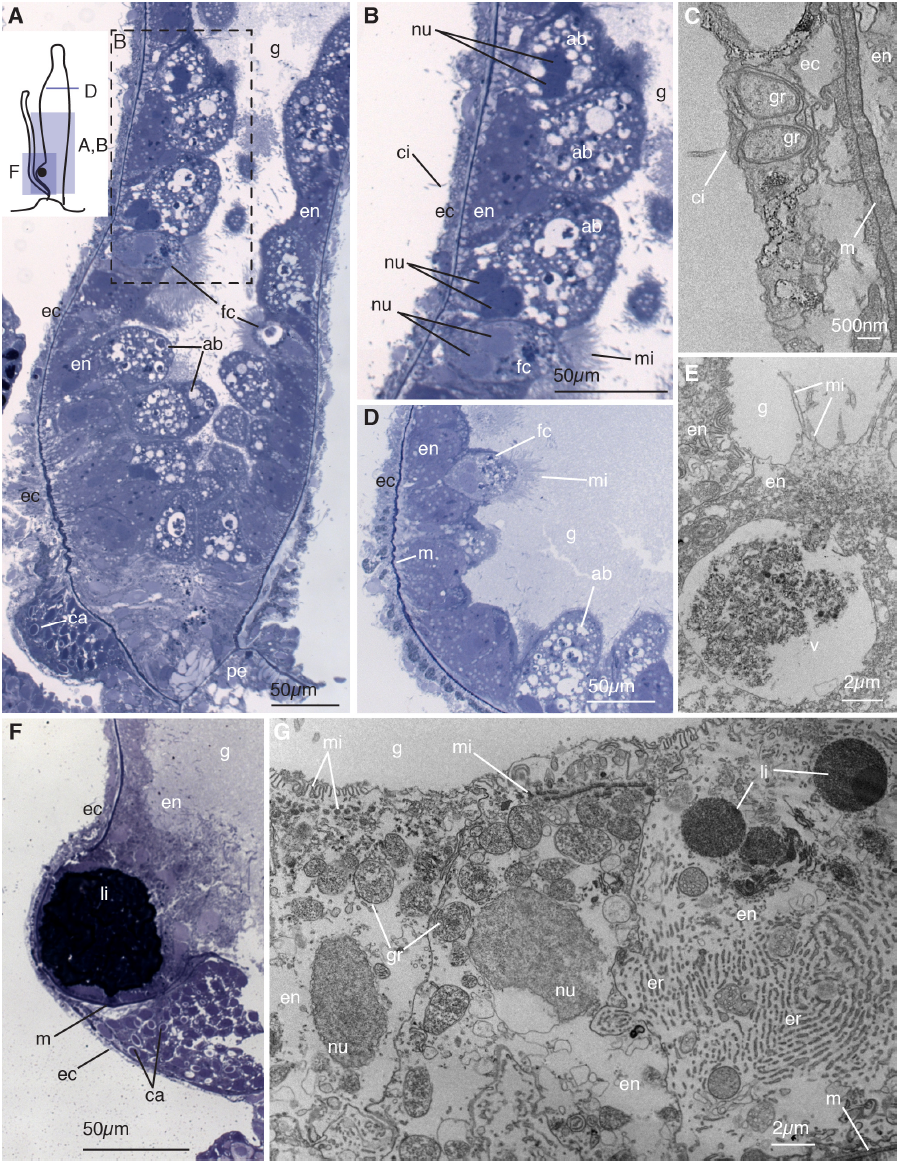
Palpon. (A) Longitudinal thick section of the base of the palpon. Schematic in the top left shows the orientation of panels A-B, D, and F. Visible are funnel cells (fc) and absorptive cells (ab) in the endoderm (en) lining the gastric cavity (g), the ectoderm (ec) including the reduced basigaster region with forming capsules (ca), and the peduncle (pe) that connects to the stem. (B) Enlarged image of region boxed in B showing the absorptive and funnel cells with microvilli (mi). Pairs of nuclei (nu) within a single cell are visible. Ectodermal cilia (ci) are also visible. (C) Longitudinal electron micrograph of the ectoderm showing cilia and beta granules (gr), and mesoglea (m). (D) Transverse thick section of the mid-region of the palpon, showing funnel cells endodermally and thin ectoderm. (E) Longitudinal electron micrograph of the endoderm, showing funnel cells with microvilli arranged in clusters at the apex, containing a large vacuole (v). (F) Longitudinal thick section of the base of the palpon, showing the mesoglea (m) partially surrounding a lipid droplet (li). A reduced basigaster can be observed basal to the droplet. (G) Longitudinal electron micrograph of endoderm cells from surface to mesoglea, showing rows of microvilli, a well developed endoplasmic reticulum (er), beta granules, and lipid droplets. Panels A-B, D, and F are thick sections (0.5-0.75 μm) stained with toluidine blue, and panels C, E, and G are electron micrographs (90nm).

The ectoderm of the palpon contains small cells which are occasionally ciliated (ci, Fig. 4B, C). These cells are thin and contain multiple beta granules (gr, Fig. 4C) as defined by Fawcett (1967). Developing nematocysts are only found in a region near the base of the palpon (ca, Fig. 4A, F). This region contains capsules at multiple stages of development similar to those observed in the gastrozooid basigaster.

Some palpons contain a large droplet near the base of the palpon, which creates a pronounced protrusion of the body. This droplet resides within the endodermal tissue, adjacent and apically to the site of nematocysts formation (li, Fig. 4F). The droplet shows high affinity to the toluidine blue stain used here, suggesting that it is composed of lipids. Within the endodermal cells, small, homogeneous droplets can be observed which are also likely lipids (li, Fig. 4G). The presence of oil droplets has been used to distinguish *Nanomia cara* from *Nanomia bijuga* (Totton, 1954). Our findings for *Nanomia bijuga* suggests this character cannot be used as a distinguishing feature. The lipid droplet could have multiple functions, including nutrient storage, buoyancy, or accumulation of compounds that deter predation.

### Male Gonodendra

#### Background

The compound sexual reproductive structures found in siphonophores are called gonodendra (Totton, 1965). *Nanomia bijuga* has male and female gonodendra each with with multiple gonophores (Carré, 1969). The gonodendra are inserted close to the base of the palpon, with male and female gonodendra alternating positions left and right from palpon to palpon (Dunn and Wagner, 2006). On the male gonodendron, individual gonophores are borne on a stalk connected to the base of the palpon (Carré, 1969). The gonophores are hypothesized to be greatly reduced medusae (Totton, 1965). The male gonophores of *Nanomia bijuga* do not maintain an umbrella in contrast to other closely related species of siphonophore (Carré, 1969). The male gonophore contains a blind internal cavity known as the spadix, and a saccular ectoderm (Daniel, 1985).

#### Results

The male gonophores examined here contained a large population of ectodermal sperm progenitor cells (sp, Figs. 5A-C). Sperm progenitor cells are located beneath a thin monolayer of ciliated ectodermal cells, forming an envelope around the saccular structure (ec, Figs. 5A-C). While it is unclear whether sperm maintain a connection to the underlying mesoglea, the ectodermal cells appear to be connected only to one another and not to the basement membrane (ec, Figs. 5A-C). The nuclei of sperm progenitor cells in small gonophores were round and filled the majority of the cytoplasm, while those in larger gonophores were elongated slightly (sp, Figs. 5A-C). Sperm progenitor cells were found at uniform stages of development in the gonophores (sp, Figs. 5A-C). No feeding structures were visible within the spadix of the gonophores, which approached but did not reach the surface of the gonophore in any specimen observed (Fig. 5A, C). The gastric cavity of the spadix is lined with compact, endodermal cells containing small granules (en, Fig. 5A, C).

**Figure 5.**
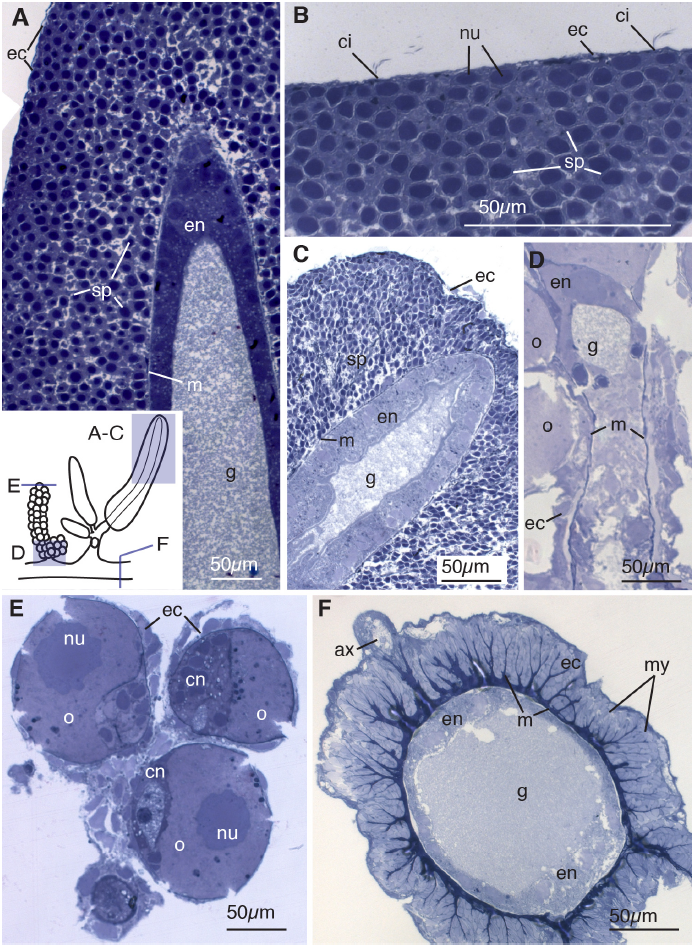
Gonodendra and Stem. (A) Longitudinal thick section of the ectoderm of the male gonophore. Schematic in the bottom right shows orientation of panels A-F. Sperm progenitor cells (sp) are located between a thin layer of ectodermal cells (ec) and the mesoglea (m). Compact endodermal cells (en) line the gastric cavity (g) of the spadix. (B) Longitudinal thick section of the surface of a male gonophore, showing cilia (ci) at regular intervals on thin ectodermal cells. Beneath the ectodermal cells are sperm progenitor cells with nuclei (nu) that fill most of the cellular space. (C) Longitudinal section showing the tip of a large male gonophore. The spadix reaches nearly the surface of the ectoderm but no opening is visible. The nuclei of the sperm progenitor cells of the large gonophore are elongated toward the ectodermal surface. (D) Longitudinal section of the female gonodendron stalk showing the tissue organization of the oocytes (o) which form ectodermally. (E) Transverse section of young oocytes showing endodermal canal cells (cn) and large nuclei. (F) Transverse section of the siphosomal stem, showing radial extentions of mesoglea lined with ectodermal cells containing myofibrils (my). The large axon (ax) is dorsal. Endodermal cells (en) are compact and thickened at the apical and dorsal sides of the gastric cavity (g). All panels are thick sections (0.5-0.75 μm) stained with toluidine blue.

### Female Gonodendra

#### Background

A thorough account of the the female gonodendron has been given by Carre (1969) as follows. The female gonodendron is a compound structure composed of a stalk and gonophores. The stalk of the gonodendron contains an internal cavity that connects to the gastric cavity near the palpon base. Individual gonophores at various stages of development are connected to the gonodendron stalk by a pedicel. Each gonophore is a reduced medusa and contains a single oocyte of ectodermal origin.The cytoplasm of mature oocytes is spotted with large vacuoles filled with vitellus. Two large looped gastrodermal canals connect to the peduncle and wrap around the oocyte .

#### Results

The stalk of a young female gonodendron, similar to that of the male gonophore, contains compact endodermal cells surrounding a tubular gastric cavity (en and g, Fig. 5D). Developing oocytes reside in between the mesoglea and ectodermal cells (ec, Figs. 5D-E). The mesoglea of the gonodendron is thin near the site of oocyte connection (m, Fig. 5D). In young oocytes the nucleus occupies nearly half of the cytoplasm (nu, Fig. 5E) and endodermal canals can be observed (cn, Fig. 5E).

### Siphosomal Stem

#### Background

All zooids of a colony are attached to the stem of the colony (Totton, 1965). The gastric cavity of the stem and the gastric cavities of each of the zooids are continuous (Totton, 1965). The stem of *N. bijuga* is highly contractile (Mackie, 1973). Mackie (1978) described the epidermis of the stem, focusing on the presence of myofibrils lining the mesoglea as well as the axons cells present at the dorsal midline.

#### Results

The myofibrils, as identified by Mackie (1978), are located along the stem are arranged in tightly packed columns along radial extensions of the mesoglea (my, Fig. 5F). These myofibrils are contained within ectodermal cells (ec, Fig. 5F). The gastrodermis of the stem is made up of small epithelial cells which are thickened on the dorsal and ventral sides of the gastric cavity (en, Fig. 5F). At the dorsal ridge of the stem are the cells of the giant axon, which are enclosed by a monolayer of ectodermal cells (ax, Fig. 5F).

### Neural Cells

#### Background

Cnidarians show significant density variation in neuronal aggregation along the body axis, contrary to the initial expectation of a diffuse nerve net (Pantin, 1952). Many species show aggregations in the form of nerve rings (Spencer and Satterlie, 1980) as observed in the buccal region of hydrozoans (Passano, 1963; Kinnamon and Westfall, 1981; Koizumi et al., 1992). Immunohistochemistry studies in siphonophores indicate a complex nervous system with nerve rings and collars in addition to a diffuse nerve net (Grimmelikhuijzen et al., 1986).

*Rfamide* encodes a neuropeptide (Plickert et al., 2003) and is used here as an indicator of a sub-population of mature nerve cells. It is a well-studied gene family in all major groups of cnidarians (Grimmelikhuijzen, 1985; Grimmelikhuijzen et al., 1988, 1991; Plickert, 1989; Koizumi et al., 1992; Moosler et al., 1996; Darmer et al., 1998; Mitgutsch et al., 1999; Anderson et al., 2004; Watanabe et al., 2009).

A previous immunohistochemistry study using an RFamide antibody performed by Grimmelikhuijzen et al (1986) revealed the following structures as RFamide positive: scattered ectodermal cells in gastrozooids, tentacles, tentilla and the pneumatophore, nerve rings at the base of the pneumatophore, gastrozooid, palpons, and gonozooids, transverse bands along the stem located one per cormidium, also known as collars, and cells of the giant axon within the stem. No previous *in situ* hybridization studies have been conducted on neuronal genes in siphonophores.

#### Results

We compared the NB*-*RFamide sequence to neuropeptides of this protein family described in other cnidarians (Fig. 6A), as summarized by Grimmelikhuijzen et al. (2004). The NB*-*RFamide shown is most similar to Pol-RFamide-II which is the RFamide of *Polyorchis penicillatus* used as a query (Fig. 6A). We characterized the expression of *nb-rfamide* by *in situ* hybridization. The expression patterns obtained here correspond very well with immunohistochemistry studies by Grimmelkhuijzen et al. (1986). The pneumatophore shows scattered *nb-rfamide* positive neurons, which are clustered near the apex and less dense near the base (ap, Fig. 6B). These cells are most dense in a ring around the apical pore.

**Figure 6.**
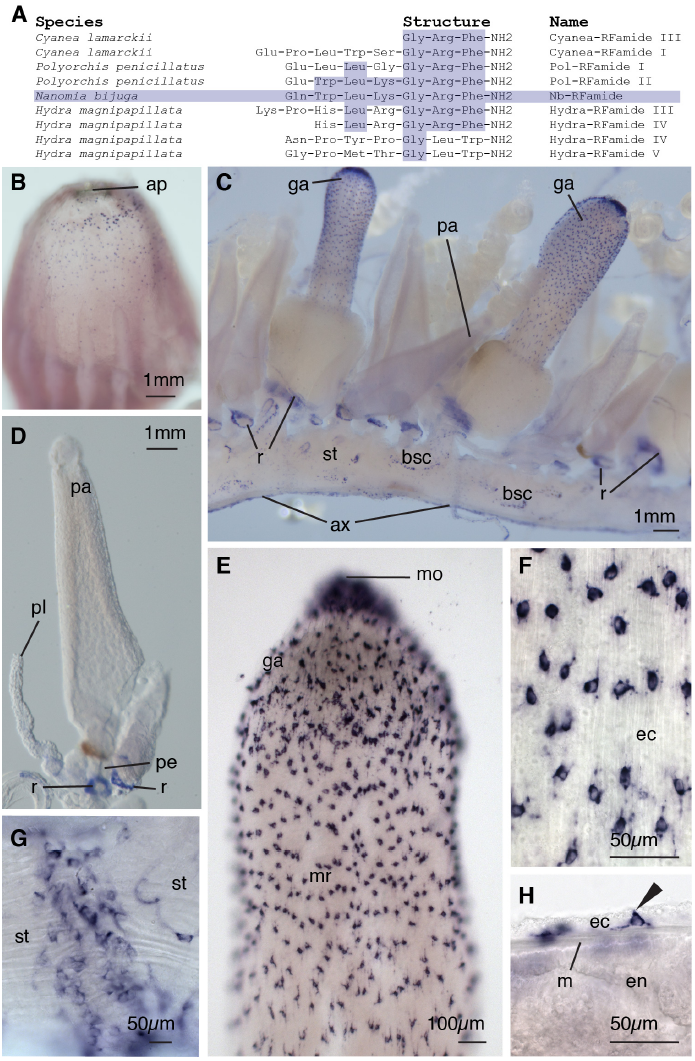
Nerve Cells. In *situ nb-rfamide* hybridization. (A) Table showing amino acid structures of selected cnidarian neuropeptides, modified from Grimmelikhuijzen (2004). Amino acids shared with Nb- RFamide are shaded. (B) The pneumatophore with the anterior facing upward, showing apical pore (ap). *nb-rfamide* positive cells are scattered most densely in a ring posterior to the apical pore, and less densely toward the mid region of the pneumatophore. (C) The cormidia of the siphosome, anterior to the left, with gastrozooids (ga) and palpons (pa) ventrally attached to the stem (st), and the giant axon (ax) on the dorsal side. Bracteal scars (bsc) are visible. *nb-rfamide* expression can be observed in rings (r) around palpon and gastrozooid peduncles above their point of attachment on the siphosomal stems. (D) Palpon showing rings of *nb-rfamide positive* cells at the base of the peduncle (pe) and at the base of the palpacle (pl). No *nb-rfamide positive* cells are observed in the palpon body or palpacle. (E) Gastrozooid hypostome showing ectodermal *nb-rfamide* positive cells with higher densities near the mouth (mo) and lower density in the mid-region (mr) of the hypostome. (F) Enlarged image of ectodermal (ec) neuronal cells in the hypostome of the gastrozooid. (G) Whole mount of the siphosomal stem showing transverse collars of *nb-rfamide* positive cells. (H) Whole mount of ectoderm, mesoglea (m), and endoderm (en) in the hypostome of the gastrozooid. *nb-rfamide* signal is absent in the endoderm. Morphology of ectodermal *nb-rfamide* positive cell (arrow) suggests sensory function.

The siphosomal stem shows scattered *nb-rfamide* positive neurons (st, Fig. 6C), possibly corresponding to bracteal scars (bsc, Fig. 6C). The giant axon is visible along the ventral side of the nectosome and the dorsal side of the siphosome (ax, Fig. 6C), opposite to the position of their respective zooids. The position of the nectosomal zooids relative to the siphosomal zooids has been identified as a shared feature of representatives from the family Agalmatidae, to which *Nanomia bijuga* belongs (Dunn et al., 2005). Members of this group have the nectophores attached dorsally, whereas zooids of the siphosome are attached ventrally (Dunn et al., 2005). The shift in orientation of the giant axon observed in this study suggests a 180° torsion between these two regions early during development, as hypothesized by Dunn (2005). Under high magnification there are transverse bands or collars on the stem located posterior to each gastrozooid (Fig. 6G). Collars are likely associated with regions of circular musculature where the stem can be constricted, and are located posterior to the gastrozooids (Grimmelikhuijzen et al., 1986).

Gastrozooids have large numbers of scattered *nb-famide* positive cells in the hypostome (ga, Fig. 6B, E), with the highest density in the area surrounding the mouth. These cells were only observed in the ectoderm (ec, Fig. 6H) and the cell morphology suggests a sensory function (arrow, Fig. 6H). Rings of *nb-rfamide* positive cells are present at the base of each gastrozooid (Fig. 6C).

The palpons also showed a dense ring of *nb-rfamide* positive neurons at their base, near the junction with the stem (pa, Fig. 6D). No patterning of scattered cells was observed in the palpon, indicating the palpon does not function primarily as a sensory structure. The palpon lacked any *nb-rfamide* signal near the tip of the palpon in the region homologous to the mouth of the gastrozooid.

## Conclusion

The distribution of cellular morphologies furthers understanding about zooid structure and function. For example, multiple hypotheses have been provided for the function of palpons, including that they act as tactile or excretory structures (Leuckart, 1853; Mackie et al., 1988). The microanatomy of the palpon, as observed in this study, supports the hypothesis that the palpon is primarily involved in digestion (Mackie et al., 1988). The endoderm of the palpon is packed with specialized cells possessing structures for particle capture, such as the funnel cells and absorptive cells. The presence of lipid droplets also suggests a possible role in long-term energy storage. The lack of *nb-rfamide* positive neurons near the tip suggests that the palpon of *Nanomia bijuga* is not primarily a sensory zooid. A deeper understanding of the microanatomy also reveals new questions for further investigation. For example, the function of the di-nucleate cells as well as relatively well developed endodermal endoplasmic reticula in the palpon remains unknown.

The microanatomy of the pneumatophore also provides a context for a discussion of its function. Pneumatophores are often cited as providing buoyancy for the colony (Pickwell et al., 1964), but they are in many species of siphonophore small relative to the total size of the organism, which makes this function unlikely (Jacobs, 1937). Our findings indicate that the pneumatophore has sensory function. The gas-filled structure may be used to detect orientation or depth. In the case of *Nanomia bijuga*, the development of the tissue layers in this structure results in the gas chamber being surrounded by three ectodermal and two endodermal tissue layers as well as mesoglea, gastrovascular space, and chitin. These multiple layers may serve to prevent gas diffusion. The presence of an apical pore suggests that the overall pressure and buoyancy of the structure can be regulated. The large population of gas producing cells within the gas chamber is evidence that the float is a center of dynamic cellular processing in a mature colony. The pneumatophore and the production of gas in the siphonophore colony merits further functional investigation.

Throughout the *Nanomia bijuga* colony there are regions of both endodermal and ectodermal specialization, including tissues with unique morphology for a singular function as well as tissues containing a diversity of cell types. The cells in the pneumatophore derived from invaginated ectoderm are highly specialized. Part of these ectodermally derived cells remain in a tight mono-epithelial layer and, unlike any other cells in the siphonophore colony, are involved in chitin production. The aeriform population, derived from the same tissue during development, is not arranged in an mono-epithelial layer and produces gas. These two cell populations are formed from the same unique event in siphonophore development, yet in a mature structure have different function and tissue morphology, both of which are unique from any other region in the colony. In the hypostome, the region of the gastrozooid adjacent to its mouth, the endoderm is many times thicker than the ectoderm. Multiple endodermal cell types are found in this region of highly expandable tissue, including three gland cell types. Adjacent to the hypostome, in the basigaster, is an area of ectodermal specialization where nematocysts are forming *en masse*. Both these regions, the hypostome and basigaster, are located within a single zooid type, indicating that function is not only achieved through zooid specialization, but also through regional tissue specialization of both gastrodermis and epidermis.

It is commonly asserted that cnidarians are simple animals composed entirely of an extracellular matrix sandwiched between two single-layered epithelia. The results presented here reinforce that this is clearly not the case. The aeriform tissue and gastrozooid basigaster ectoderm both include cells that are not in direct contact with the mesoglea, and therefore are not mono-layered epithelia. Our data suggests it is unlikely that these tissues are pseudostratified mono-layers, where the relative position of the nuclei can create the appearance of multiple epithelial layers within a single section. Within these two regions, cell boundaries and nuclei are visible within the same plane; furthermore cells in these regions are spherical, and there are no visible cellular projections extending to connect with mesoglea. This is in contrast to regions such as the endoderm of the hypostome, where nuclei are positioned at a distance from the mesoglea, but cellular connections with the basement membrane are visible, indicating that this region is a thick mono-epithelial layer.

## Acknowledgements

Thanks to Steven H.D. Haddock for making collection of *Nanomia bijuga* possible, and for providing valuable commentary to this manuscript. Thanks to Sophia Tintori for initial protocol development and technical support. Thanks to the Leduc Bioimaging Facility at Brown University, as well as Geoffrey Williams for excellent technical support with microscopy. Thanks to Catriona Munro and Phil R. Pugh for their helpful comments with respect to siphonophore literature and biology. This work was supported by Alan T. Waterman award from the National Science Foundation, NSF grant DEB-1256695, and by the EPSCoR Summer Undergraduate Research Fellowship program.

## Literature Cited

Anderson PA, Thompson LF, Moneypenny CG. 2004. Evidence for a common pattern of peptidergic innervation of cnidocytes. The Biological Bulletin 207:141–146.

Bardi J, Marques AC. 2007. Taxonomic redescription of the Portuguese man-of-war, Physalia physalis (Cnidaria, Hydrozoa, Siphonophorae, Cystonectae) from Brazil. Iheringia Série Zoologia 97:425–433.

Bosch TC, David CN. 1987. Stem cells of Hydra magnipapillata can differentiate into somatic cells and germ line cells. Developmental Biology 121:182–191.

Campbell RD. 1967. Cell proliferation and morphological patterns in the hydroids *Tubularia* and Hydractinia. Journal of embryology and experimental morphology 17:607–616.

Carré D, Carré C. 1995. Ordre des Siphonophores. In: Doumenc D (ed) Traité de Zoologies: Anatomia, Systematique, Biologie. Vol. 3(2). Paris: Masson. p 523–596.

Carré D. 1969. Étude histologique du développement de Nanomia bijuga (Chiaje, 1841), Siphonophore Physonecte, Agalmidae. Cahiers de Biologie Marine 10:325–341.

Carré D. 1971. Étude du développement d’ Halistemma rubrum (Vogt, 1852), Siphonophore Physonecte, Agalmidae. Cahiers de Biologie Marine 12:77–93.

Chun C. 1891. Die Canarischen Siphonophoren I. Stephanophyes superba und die Familie der Staphanophyiden. Abhandlungen hrsg von der Senckenbergischen Naturforschenden Gesellschaft 16:553–627.

Claus C. 1878. Über *Halistemma tergestinum* n. sp. nebst Bemerkungen über den feinern Bau der Physophoriden. Arbeiten aus dem Zoologischen Instituen der Universität Wien und der Zoologischen Station in Triest 2:199–202.

Daniel R. 1985. The fuana of India and the adjacent countries. Coelenterata: Hydrozoa, Siphonophora. Publication of the Zoological Survey of India:440.

Darmer D, Hauser F, Nothacker H, Bosch T, Williamson M, Grimmelikhuijzen C. 1998. Three different prohormones yield a variety of Hydra-RFamide (Arg-Phe-NH2) neuropeptides in Hydra magnipapillata. Biochem J 332:403–412.

Dunn CW, Howison M, Zapata F. 2013. Agalma: an automated phylogenomics workflow. BMC bioinformatics 14:330.

Dunn CW, Pugh PR, Haddock SH. 2005. Molecular phylogenetics of the Siphonophora (Cnidaria), with implications for the evolution of functional specialization. Systematic Biology 54:916–935.

Dunn CW, Wagner GP. 2006. The evolution of colony-level development in the Siphonophora (Cnidaria: Hydrozoa). Development genes and evolution 216:743–754.

Fawcett DW. 1967. An Atlas of Fine Structure: the cell, its organelles and inclusions. Philadelphia and London: W. B. Saunders Company.

Garstang W. 1946. The morphology and relations of the Siphonophora. Quarterly Journal of Miscroscopical Science 87:103–193.

Genikhovich G, Technau U. 2009. In situ hybridization of starlet sea anemone (Nematostella vectensis) embryos, larvae, and polyps. Cold Spring Harbor protocols 2009:pdb–prot5282.

Grimmelikhuijzen C, Graff D, Koizumi O, Westfall JA, McFarlane ID. 1991. Neuropeptides in coelenterates: a review. In: Coelenterate Biology: Recent Research on Cnidaria and Ctenophora. Springer. p 555–563.

Grimmelikhuijzen C, Hahn M, Rinehart KL, Spencer AN. 1988. Isolation of 2(Pol-RFamide), a novel neuropeptide from hydromedusae. Brain research 475:198–203.

Grimmelikhuijzen C, Spencer AN, Carré D. 1986. Organization of the nervous system of physonectid siphonophores. Cell and tissue research 246:463–479.

Grimmelikhuijzen C. 1985. Antisera to the sequence Arg-Phe-amide visualize neuronal centralization in hydroid polyps. Cell and tissue research 241:171–182.

Grimmelikhuijzen CJ, Williamson M, Hansen GN. 2004. Neuropeptides in cnidarians. Cell Signalling in Prokaryotes and Lower Metazoa. Springer. p 115–139.

Haddock SH, Dunn CW, Pugh PR. 2005. A re-examination of siphonophore terminology and morphology, applied to the description of two new prayine species with remarkable bio-optical properties. Journal of the Marine Biological Association of the United Kingdom 85:695–707.

Jacobs W. 1937. Beobachtungen über das Schweben der Siphonophoren. Zeitschrift für Vergleichende Physiologie 24:583–601.

Johnson M, Zaretskaya I, Raytselis Y, Merezhuk Y, McGinnis S, Madden TL. 2008. NCBI BLAST: a better web interface. Nucleic acids research 36:W5–W9.

Kinnamon JC, Westfall JA. 1981. A three dimensional serial reconstruction of neuronal distributions in the hypostome of a Hydra. Journal of Morphology 168:321–329.

Koizumi O, Itazawa M, Mizumoto H, Minobe S, Javois LC, Grimmelikhuijzen CJ, Bode HR. 1992. Nerve ring of the hypostome in hydra. I. Its structure, development, and maintenance. Journal of Comparative Neurology 326:7–21.

Leloup E. 1935. Les siphonophores de la Rade de Villefranche-sur-Mer (Alpes, Maritimes, France). Bulletin de Musée Royal d’Histoire Naturelle de Belgique 11.

Leuckart R. 1853. Zoologische Untersuchungen. I. Die Siphonophoren. Giessen: J. Ricker’sche Buchhandlung.

Mackie GO, Pugh PR, Purcell JE. 1988. Siphonophore biology. Advances in Marine Biology 24:97–262.

Mackie GO. 1973. Report of giant nerve fibres in Nanomia. Publications of the Seto Marine Biological Laboratory 20:745–756.

Mackie GO. 1978. Coordination in physonectid siphonophores. Marine & Freshwater Behaviour & Phy 5:325–346.

Metschnikoff E. 1870. Contributions to the knowledge of siphonophores and medusae. Mémoires de la Société des Amis des Sciences Naturelles d“Anthropologie et d”Ethnographie 8:295–370.

Mitgutsch C, Hauser F, Grimmelikhuijzen CJ. 1999. Expression and Developmental Regulation of the Hydra-RFamide and Hydra-LWamide Preprohormone Genes in Hydra: Evidence for Transient Phases of Head Formation. Developmental Biology 207:189–203.

Moosler A, Rinehart KL, Grimmelikhuijzen CJ. 1996. Isolation of Four Novel Neuropeptides, the Hydra-RFamides I-IV, from Hydra magnipapillata. Biochemical and biophysical research communications 229:596–602.

Pantin C. 1952. Croonian Lecture: The Elementary Nervous System. Proceedings of the Royal Society of London Series B, Biological Sciences: 147–168.

Passano LM. 1963. Primitive nervous systems. Proceedings of the National Academy of Sciences of the United States of America 50:306.

Pickwell GV, Barham EG, Wilton JW. 1964. Carbon monoxide production by a bathypelagic siphonophore. Science 144:860–862.

Plickert G, Frank U, Müller WA. 2012. Hydractinia, a pioneering model for stem cell biology and reprogramming somatic cells to pluripotency. International Journal of Developmental Biology 56:519.

Plickert G, Schetter E, Verhey-Van-Wijk N, Schlossherr J, Steinbuchel M, Gajewski M. 2003. The role of alpha-amidated neuropeptides in hydroid development-LWamides and metamorphosis in Hydractinia echinata. International Journal of Developmental Biology 47:439–450.

Plickert G. 1989. Proportion-altering factor (PAF) stimulates nerve cell formation in Hydractinia echinata. Cell differentiation and development 26:19–27.

Pugh PR. 1999. Siphonophorae. In: South Atlantic Zooplankton, D. Boltovskoy, ed. Leiden: Backhuys Publishers. p 467–511.

Rozen S, Skaletsky H. 1999. Primer3 on the WWW for general users and for biologist programmers. In: Bioinformatics methods and protocols. Springer. p 365–386.

Siebert S, Anton-Erxleben F, Bosch TC. 2008. Cell type complexity in the basal metazoan Hydra is maintained by both stem cell based mechanisms and transdifferentiation. Developmental Biology 313:13–24.

Siebert S, Goetz FE, Church SH, Bhattacharyya P, Zapata F, Haddock SHD, Dunn CW. 2014. Stem cells in a colonial animal with localized growth zones. bioRxiv. doi:10.1101-001685

Siebert S, Robinson MD, Tintori SC, Goetz F, Helm RR, Smith SA, Shaner N, Haddock SH, Dunn CW. 2011. Differential gene expression in the siphonophore Nanomia bijuga (Cnidaria) assessed with multiple next-generation sequencing workflows. PLoS ONE 6:e22953.

Spencer AN, Satterlie RA. 1980. Electrical and dye coupling in an identified group of neurons in a coelenterate. Journal of neurobiology 11:13–19.

Thomas MB, Edwards NC. 1991. Cnidaria: hydrozoa. In: Microscopic Anatomy of Invertebrates. Vol. 2. p 91–183.

Totton AK. 1954. Siphonophora of the Indian Ocean: Together with Systematic and Biological Notes on Related Specimens from Other Oceans. University Press.

Totton AK. 1965. A Synopsis of the Siphonophora. London: British Museum (Natural History).

Watanabe H, Fujisawa T, Holstein TW. 2009. Cnidarians and the evolutionary origin of the nervous system. Development, growth & differentiation 51:167–183.

Weismann A. 1892. Das Keimplasma. Eine Theorie der Vererbung. Gustav Fischer.

Willem V. 1894. La structure du palpon chez *Apolemia uvaria* Esch., et les phénomènes de l’absorption dans ces organes. Bulletin de l’Académie Royale de Belgique Classe des Sciences 27:354–363.

